# Testing the null model for polyandry: the need to breed explains multiple mating and constrains trading-up

**DOI:** 10.64898/2026.05.08.723703

**Authors:** Danica S McCorquodale, Jacob D Berson, Robert J Dugand, Natasha R LeBas, Joseph L Tomkins

## Abstract

In most species, unmated individuals run the risk of dying with zero fitness. This strong selection on virgin females to mate may also explain why females subsequently remate, despite fitness costs; all that is required is a genetic correlation between virgin and non-virgin mating propensity. Despite being the null model for the evolution and maintenance of polyandry, this hypothesis has received no empirical test. We performed separate quantitative genetic and artificial selection experiments to test the presence of this cross-context genetic correlation in the cow-pea weevil, *Callosobruchus maculatus*. A quantitative genetic experiment did not find evidence of the hypothesised genetic correlation. However, after 13 generations of artificial selection on virgin mating latency, we found strong evidence for evolutionary divergence in remating latency. Females from lines selected for longer virgin mating latency took approximately twice as long to remate and, were less polyandrous if their virgin mating latency was longer. There was no evidence that females could mate indiscriminately and then ‘trade-up’, rather, trading up could only occur if virgin discrimination was present. Selection against virgin death will thus constrain both the evolution of non-virgin discrimination and trading up, increasing rates of polyandry. These findings reveal a genetic correlation between virgin and non-virgin latency to mate suggesting that polyandry may be maintained because of the need to breed.

## Introduction

Polyandry has given rise to the evolution of an extraordinary diversity of adaptations to both pre- and post-copulatory sexual selection [1, 2]. One of the central questions raised by the multiple mating of females is why females would mate more often than is optimal for their fitness [1-5]. There are several adaptive hypotheses for the evolution of polyandry [6]. For example, ‘convenience polyandry’ posits that multiple mating is an adaptive consequence of reducing the costs of sexual harassment – readily mate to minimise the negative consequences of harassment [3]. Alternatively, there may be cross-sex pleiotropic constraints, where genes that adaptively increase the mating frequency of males also (maladaptively) increase mating rates in females [4, 7, 8]. One hypothesis that has received, to our knowledge, no empirical attention, oft considered the null model, is that polyandry is an evolutionary by-product of the need to mate at least once, aka the ‘wallflower effect’ [9]. Sexually mature, virgin females not only waste time not reproducing, but also run the risk of a ‘virgin death’ [5, 10], see also [11, 12]. These costs may reduce female choosiness in virgin, but also non-virgin matings. The wallflower hypothesis has important ramifications for polyandry as well as the evolution of mating preferences generally [13], because it articulates the existence of a ‘preference-dependent’ risk of virgin death [9]. Kokko and Mappes’ [5, 10] models combine the avoidance of the cost of remaining virgin with either a *plastic* response of the female, that allows adjustment of subsequent decisions (allows females to ‘trade-up’) or a *fixed* response where there is an implicit (genetic) constraint, such that the avoidance of virgin death ‘carries on to later stages in life’ [5].

The extent to which females can plastically alter their behaviour, *versus* being genetically constrained, can be considered from a quantitative genetic perspective. For example, genetic correlations arise where the expression of two different phenotypic traits depend on common genetic variation, either through linkage or pleiotropy [14]. Where this occurs, selection on one trait necessarily affects the other, and the traits are constrained in their evolution. For example, genetic correlations between life stages for the same trait can exert constraints across life stages [15]. Similarly, under the environmental threshold model of the conditional strategy where organisms make decisions based on environmental cues [16], we expect those genotypes that are more sensitive to the strength of the cue in one environment to be more sensitive to that same cue in another environment, for example, at another developmental stage, or in this case, a virgin *versus* a non-virgin mate encounter [17]. If the environmental cue is the courtship of a male, and the decision is between mating or not, the suggestion from both these theoretical standpoints is that selection that favours females that are responsive enough to male courtship to mate once will also result in them being likely to mate again, if there is a genetic correlation between the choices made at these two times in the female’s life. Thus, even though the optimal individual strategy may be to mate only once [5], the optimal evolutionary strategy is to mate more than once, because mating more readily, avoids the costs of remaining virgin.

Here, using two approaches, we test the genetic correlation between virgin and non-virgin latency to mate in the cow-pea weevil *Callosobruchus maculatus*. Polyandry is heritable in many species [18-25]; however, the relationship between virgin and non-virgin female latency to mate is largely unexplored (Dugand et al 2025). First, we conducted a half-sibling breeding design to directly estimate the additive genetic correlation between mating latencies of virgin and non-virgin females. Second, we artificially selected on virgin female latency to mate and quantified the correlated response in the latency to mate of non-virgin females and their degree of polyandry. Despite a lack of quantitative genetic evidence for additive genetic variance in mating latency (virgin or non-virgin females), we found that artificial selection on virgin latency to mate resulted in a correlated response in non-virgin latency to mate, supporting a positive genetic correlation between mating latencies across contexts. Crucially this carried over into changes in the expression of polyandry. Our results support the null model of polyandry, suggesting that polyandry may be maintained as an evolutionary byproduct of the need to breed.

## Methods

### Base population

The base population of *C. maculatus* was sourced from CSIRO Canberra and has been maintained as a large, outbred population since 2005. The population was maintained on black-eyed beans (*Vigna unguiculata*) at 28 °C in a Binder 240 L incubator under a 12:12 light:dark cycle. This population has been shown to have abundant additive genetic variation for a range of quantitative traits [26-30]. The selection experiment was conducted 2012-2013 and the half-sibling experiment in 2014.

### Pedigreed half-sib experiment

To initiate the pedigree, sires and dams were drawn from the base population by removing approximately 1,000 beans with eggs, which were placed individually into 1.7 mL microtubes. The tubes were checked regularly for adult beetle emergence, and virgin adults were separated from the bean and kept isolated in a microtube. Within a week of emergence, beetles were paired, with 76 sires each mated to three dams over three consecutive days. Following mating, females were isolated into a ventilated 150 mL plastic specimen vial with about 50 black-eyed beans and returned to the incubator. After the female’s death (1-2 weeks after mating), beans with eggs were isolated into punctured microtubes and placed in dam families in small zip-lock bags that had been perforated once with a hole-punch. These bags were checked daily for eclosion. We obtained offspring from 67 of the original 76 sires, and 131 of the original 532 dams (**Table S1**).

After offspring eclosion, virgin females from the 131 dam families were assayed for their latency to mate to males collected from a different family (*i*.*e*., males that did not share a sire or dam with the female). Mating in *C. maculatus* is confirmed when the male mounts and inserts his aedeagus inside the female. The time to mating was recorded with a stopwatch. Once a pair separated, the male was discarded, and the female was placed into a ventilated specimen jar along with approximately 50 black-eyed beans. The mated female was then left for 2 days. The female was then placed again with another random, virgin male and time to mating recorded. After copulation was complete, both adult beetles were discarded.

We continued the pedigree for a second generation, allowing us to collect additional data on mating and remating. To do so, 38 female offspring from 38 of the initial sire families were mated with 38 males from the stock population to generate 38 full-sib families (**Table S1**; **Fig. S1**). From these 38 families, we collected 75 males and 222 females and used these to construct a second full-sib half-sib design (**Fig. S1**). For each pair, we recorded the time to mate, but not the time to remate (**Table S1**). The offspring from these 222 full-sib families were assayed for both mating and remating as outlined above. Therefore, we recorded mating and remating times from a total of 142 sires and 353 full-sib families, though not all were drawn from the stock population (see **Table S2** for a summary of the relationships within the additive genetic relatedness matrix).

### Selection experiment

To initiate the selection experiment, we collected virgin male and female beetles from the base population. We randomly assigned 240 pairs and recorded their time to mating. Pairs were ranked on the time to mating, with the fastest 60 pairs randomly allocated to three ‘*fast*’ lines, and the slowest 60 randomly allocated to three ‘*slow*’ lines; each selection line had 20 males and 20 females. Females were allowed to lay communally on fresh black-eyed beans. The egg-bearing beans from these females were isolated into punctured microtubes and virgins collected upon eclosion. Thus, in the first generation, selection included both female and male latency to mate.

At each subsequent generation, 50 virgin females from each line were assayed for their time to mounting with males from the base population. To do this, a virgin stock population male was introduced to the female’s microtube and the time recorded. As soon as the male mounted the female, the time was recorded and the pair were separated with soft forceps. Ejaculate is not transferred in *C. maculatus* until after the second minute of mating [31] so our method ensured that no sperm was transferred. The male was discarded after the assay. The 10 females that were mounted fastest in the *fast* lines, and slowest in the *slow* lines, were identified. These females, that had been mounted but not mated, were then introduced to a container with 20 males from their own line and allowed to mate to produce the next generation of beetles. Selection continued in this way for 13 generations.

Following artificial selection on virgin female latency to mate, the presence of a genetic correlation between virgin and non-virgin mating latency can be tested by measuring the correlated response in non-virgin mating. At generation 13, we assayed female latency to mate to stock population males and used the fastest and slowest females to propagate their respective selection lines (which we subsequently did not continue). We then measured the latency to remate of the remaining females (to base population males). This was done for their first polyandrous opportunity to mate 5-7 days (mean = 5.7 days all lines) after their first mating. We used these values as our measure of remating latency. Females, that did not remate were not trialled again; however, those that did had an additional mating opportunity after 2 days (all lines). Note that, at generation 13 because the fastest mating females in *fast* lines, and the slowest mating females in *slow* lines, were set aside as foundresses for the next generation and were not assayed for remating latency, our estimates of remating latencies would be biased to be faster in the *slow* lines and slower in the *fast* lines. Consequently, our test of the selection regime effect on remating latency and probability is likely to be conservative. For the entire selection experiment and subsequent assays, all mating trials were performed blind to the identity of the line.

### Statistical analyses

#### Half sibling analysis

We used the *brms* R package [32] in R Version 4.4.0 [33] to estimate the genetic correlation between latency to mate during the first mating, and latency to mate during the second mating. To do so, we fit a generalised linear ‘animal model’ using an exponential likelihood with a log link. An animal model uses a relatedness matrix to estimate the additive genetic variance associated with each trait, and their genetic covariance [34]. In addition to including the relatedness matrix, we estimated the residual and dam (co)variances of the two traits, as well as the variances for the date the mating assays took place. We used the default flat priors for the population level effects, the default half student-t priors with three degrees of freedom and a scale parameter of 2.5 for the standard deviations of the group-level (‘random’) effects, and an LKJcorr prior with *eta* equal to five for the group-level correlations. This latter prior, places most of the prior probability on correlations between -0.5 and 0.5 and aided model convergence. We excluded females that did not mate during the second mating (824 females, 48% of those assayed).

Given the large number of females that were unreceptive during the second mating and therefore excluded from the above analysis, we conducted a supplementary analysis to estimate the level of additive genetic variance in female receptivity during the second mating. To do so, we modelled the binary trait of whether a female mated during the second mating assay by fitting a generalised linear animal model using a Bernoulli likelihood with a logit link. In this model we estimated the variance associated with the relatedness matrix (*i*.*e*., additive genetic variance) along with variances associated with the female’s mother (‘dam’) and the date that mating took place. We used a Normal prior with mean of zero and standard deviation of two for the model intercept, and an exponential prior with lambda equal to five for the standard deviations of the group-level effects.

#### Selection experiment

Mating latencies varied substantially across generations (**Fig. 1**). Therefore, we first removed this variation by taking the residuals of a fitted model, with seconds to mate (power-transformed, y^-0.34^) predicted by generation, where generation was modelled as a categorical fixed effect. Using the residuals from this model, we tested the fixed effects of regime (*fast versus slow*) and a generation-by-regime interaction (where generation was a continuous fixed effect) on mating latency. We retained generation as a continuous fixed effect. To account for the repeated measures for line, we modelled independent random slope (across generation) and intercept effects, ensuring we had the correct denominator degrees of freedom [35]. The generation-by-regime interaction provides the most robust test of a response to artificial selection, where divergence between selected traits should increase through time.

**Fig. 1.**
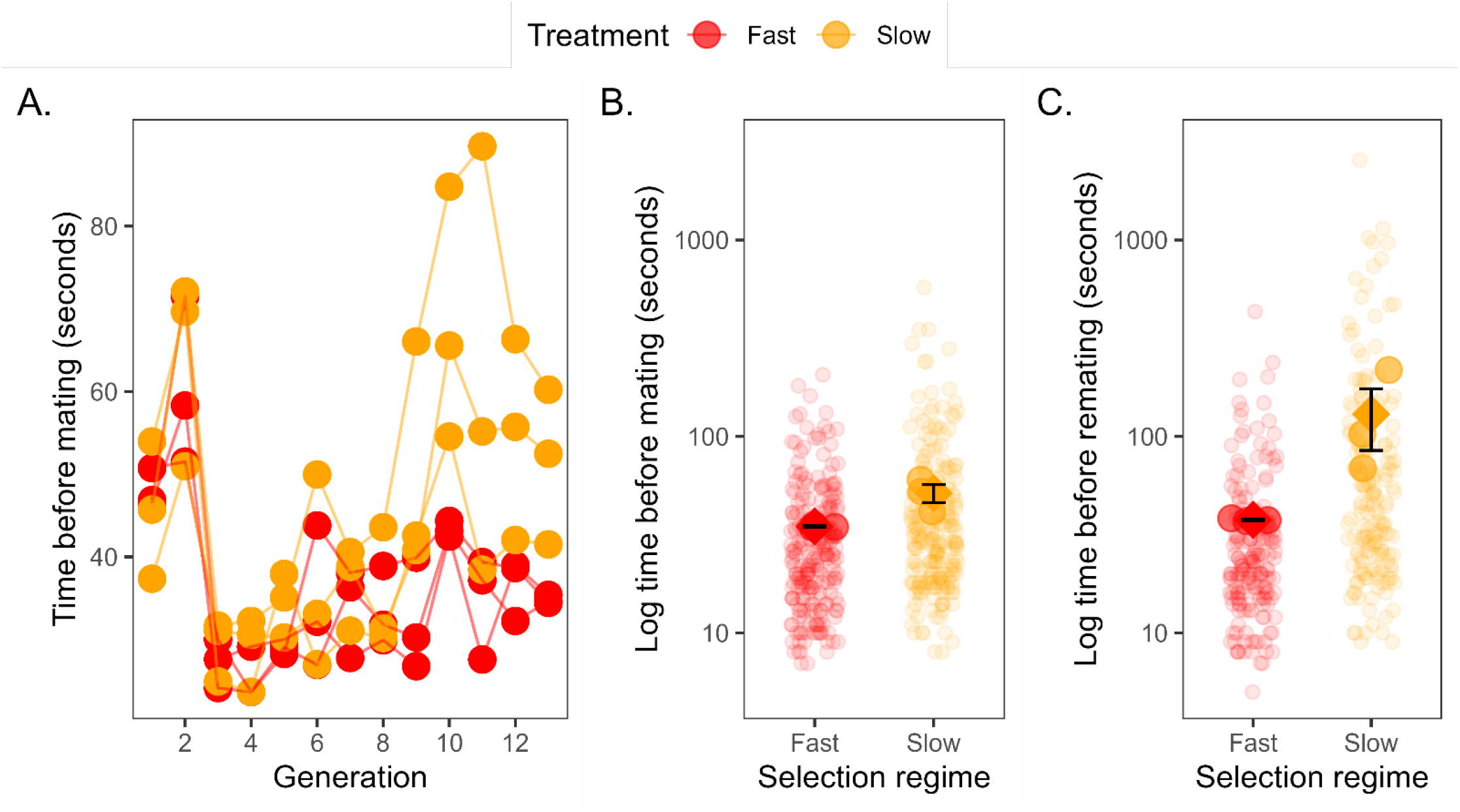
Effect of selection regime on time to (re)mating. **A**. Mean time to mate for each selection line per generation. **B**. Marginal means for the time before mating in the first mating assay. **C**. Marginal means for the time before mating in the remating assay. Slow-selected lines are shown in orange, and fast-selected lines are shown in red. In B and C, error bars are 95% compatibility intervals, the jittered transparent points are the raw data, the densely coloured centre-aligned points are line means (also note the log-scale).

Additionally, we used mating latency data from the final generation to test the effect of selection regime on the power-transformed (y^-0.30^) latency to mate data. Line was included as a random effect. To test whether there was a correlated response in remating latency, we tested the fixed effect of selection regime on the power-transformed (y^-0.34^) latency to mate data. Line was included as a random effect.

We also separately tested an interaction model, to specifically test the hypothesis that latency (y^-0.34^) to mate differentially affected latency (y^-0.34^) to remate, dependent on the selection history; selection history and mating (virgin or non-virgin) were fixed effects and we included ID as an additional random effect in this model, alongside the random effect of line.

The genetic correlation between virgin and non-virgin mating responsiveness can also be estimated by Pearson’s correlation coefficient of raw line means of mating latency for the first and second mating. In this analysis, each female was measured for both their first and second mating means in these correlations, potentially inflating the estimate with within individual effects. The mean may also be a biased measure of central tendency in these right-skewed data. To estimate a correlation, free from these two effects, we randomly assigned females either to contribute to the latency to mate median estimation or the latency to remate median estimation and then calculated the correlation of line medians repeatedly (1,000 reps) and report the median of these correlations, along with 95% sampling intervals.

Finally, we tested the relationship between the number of times out of two opportunities a female remated (polyandry), and the time a female took to mate as a virgin. To do so, we employed cumulative ordinal logistic mixed model, with a logit link function (using the package *ordinal*), where remating 0, 1, or 2 times was the dependent variable and latency to mate as a virgin was the independent variable and treatment was a fixed factor and line was a random effect. We tested the significance of the interaction with a likelihood ratio test (LRT) between the full model and the model excluding the interaction term. The proportional odds assumption was tested for by comparing the consistency of two separate logistic regressions between 0 to >1 mating and 1 mating to 2 matings.

Prior to analysis, we excluded 41 (10.5%) females that did not mate (17 from fast lines and 24 from slow lines).

All R code used to perform that analyses can be found in supplementary material.

## Results

### Half sibling analysis

Females mated more readily as virgins (estimated average latency to mate = 41 seconds, 95% compatibility intervals, CI: 32 – 52 seconds) than non-virgins (estimated average latency to mate = 213 seconds, 95% CI: 111 – 412 seconds). The estimated genetic correlation between the latency to mate during the first mating, and the latency to mate during the second mating was -0.09 (95% CI: -0.64 – 0.52, **Extended Data Fig. 2**). Given that genetic correlations ranging from -0.64 to 0.52 were compatible with our data given our model, we interpret this result as statistically unclear [36] and do not draw any inference about the presence or absence of a genetic correlation. Complete model results are available in **Extended Data Fig. 2** and **Extended Data Table 3**.

We did not find statistically clear evidence for the presence of additive genetic variance in female receptivity during the second mating, with additive genetic variance estimated to represent 9% of the total variance (95% CI: 0 – 48%). There was clear evidence that variation in female receptivity was associated with the identity of the female’s mother (39% of the variation, 95% CI: 4 – 69%) and the date the mating assay took place (52% of the variation, 95% CI: 25 – 83%). Full results are shown in **Extended Data Fig. 3** and **Extended Data Table 4**.

### Selection experiment

We found weak support for a generation-by-regime interaction effect (χ^2^ = 3.25, d.f. = 1, *P* = 0.071; see **Extended Data Table 5** for full results), where regimes became increasingly divergent through time, and in the direction in which they were selected (**Fig. 1A**). At the final generation of selection, females from *fast* lines mated significantly faster than females from *slow* lines (χ^2^ = 4.96, d.f. = 1, *P* = 0.026), although there was some overlap in 95% compatibility intervals (estimated marginal means: fast-selected lines = 25 seconds, 95% CI: 20 – 32 seconds; slow-selected lines = 33 seconds, 95% CI: 26 – 43 seconds, **Fig. 1B**).

There was a highly significant selection regime effect on female remating latency at generation 13 (χ^2^ = 24.05, d.f. = 1, *P* < 0.001) where females from *slow* lines had longer remating latencies (estimated marginal means: fast-selected lines = 24 seconds, 95% CI: 19 – 30 seconds; slow-selected lines = 47 seconds, 95% CI: 35 – 64 seconds, **Fig. 1C**). These effects are supported further by the interaction model, where there was an overall treatment effect on latency (to mate or remate) (F1,4,=11.02, P=0.029), a significant increase in the latency to remate compared to the first mating (F1,4,=8.17, P=0.046), and a significant treatment by latency (to mate or remate) interaction (F1,4,=16.76, P=0.015) revealing that lines selected for slow matings were even more slow to remate than lines selected to mate rapidly.

There was a significant Pearson’s correlation among the line means between the time to mate as a virgin and the time to remate as a non-virgin (Fig. 2; r = 0.95, 95% CI 0.61-0.99, t = 6.10, d.f. = 4, p = 0.004). This correlation across raw line means was supported by correlation across line means derived from box-cox transformed data (r= 0.93, 95% CI 0.5-0.99). Each female contributed to both the first and second mating means in these correlations, potentially inflating the estimate with within individual effects and these data are skewed. By randomly assigning females either to contribute to the latency to mate median estimation or the latency to remate median estimation, and then calculating the correlation of line medians, the median correlation coefficient was still high, r = 0.78 (Interquartile range = 0.64 - 0.87).

**Fig. 2.**
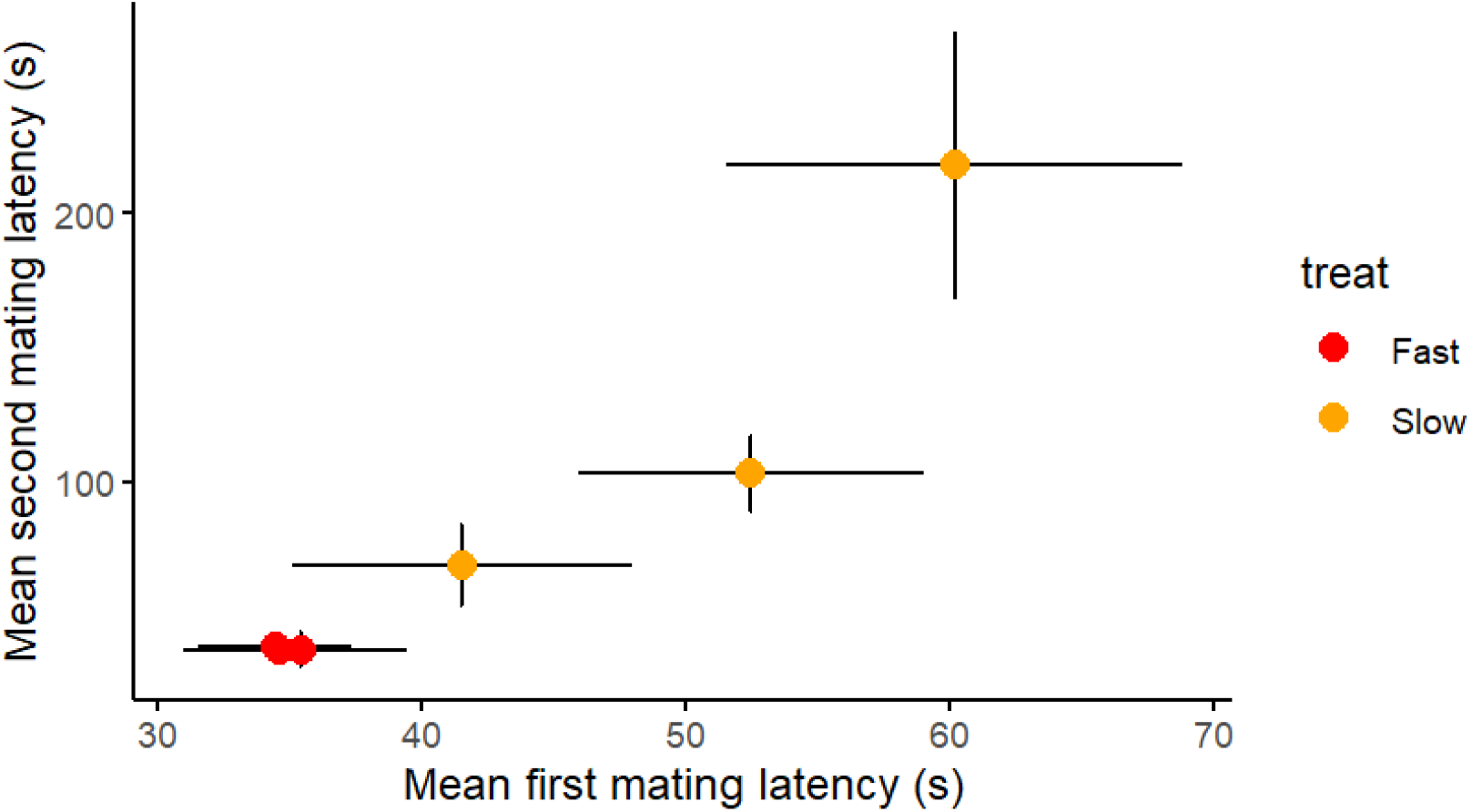
Correlation of raw line means +/-SE for the six lines selected for divergent time to first mating, and their correlated response in the time to remating.

When females were repeatedly tested for their propensity to remate, after their first mating, we found a significant interaction between the time that they took to mate as a virgin and their level of polyandry (LRT between the full model and the model without the interaction; LR=5.367, d.f.= 1, P=0.021). The interaction reveals that females with a selection-history of slow virgin matings, are less likely to remate when presented with males across two consecutive occasions, if their time to the first mating was longer; but that this was not the case in the lines with a selection history of mating rapidly (Fig. 3). The interaction remains significant when the data are limited to only first mating latencies that overlap across treatments. The proportional odds assumption was met: separate logistic regressions for the thresholds 0 to 1+2 and 0+1 to 2 have the same rank order and magnitude of effect sizes, treatment:time to first mating > treatment > time to first mating and interaction terms are consistent, P=0.09 and P=0.04. The correlation between line medians of time to first mating and mean number of remating is underpowered but consistent (t_4_ = -1.7, p = 0.162, r = -0.65).

**Fig. 3.**
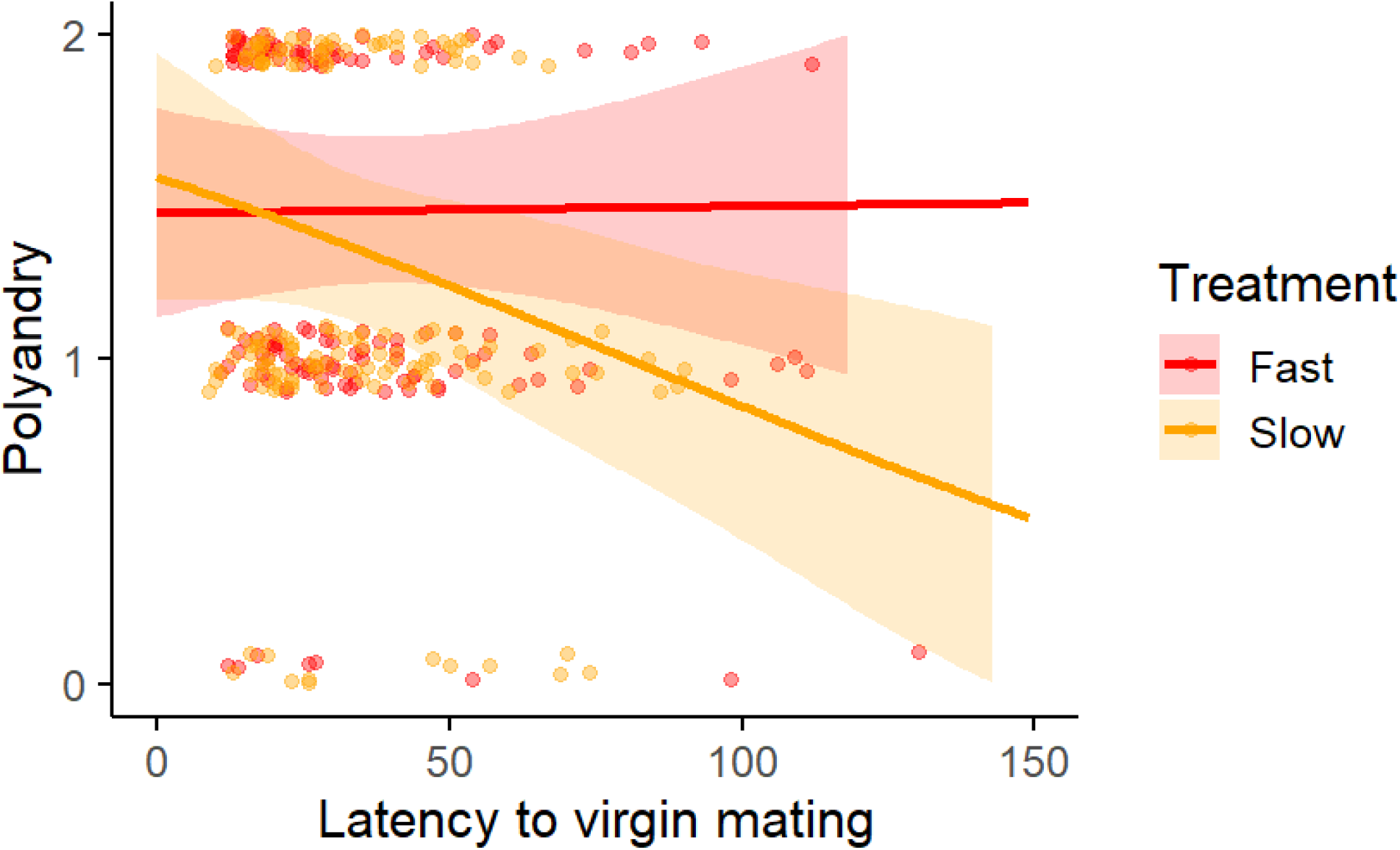
Relationship between polyandry measured as the number of times a mated female mated again in two consecutive trials, plotted against that female’s latency to mate as a virgin. The lines are predicted from the cumulative logistic mixed model; data points are jittered. Females with a selection history that increased their latency to mate as virgins, were less polyandrous when they took longer to mate as virgins. Three outlying data points are missing from the figure but not the analysis (Fast,2 females x 2 matings (161,2) (181,2), 1 female x 1 mating (206.1))

## Discussion

We aimed to test the hypothesis that the evolutionary maintenance of apparently costly polyandrous matings might arise through selection on females against virgin death [5]. The costs of remaining virgin (lost time and virgin death) are contingent upon the strength of female preferences for males, as such the costs are preference-dependent because overly choosy females risk a virgin death. Hence, the hypothesis predicts constraints in the form of a positive genetic correlation between the readiness of a female to mate as a virgin and the likelihood that she will later remate as a non-virgin [5, 9, 10]. Our results provide weak statistical support for divergence in virgin mating latency over 13 generations of selection, but strong and significant divergence in non-virgin female mating latency. This is compelling evidence for a correlated evolutionary response to selection on virgins driven by an underlying positive genetic correlation between virgin and non-virgin latency to mate. Critically we show that selection for extended latency to virgin mating, reveals a negative relationship between latency to mate as a virgin and polyandry. Our results therefore support the Wallflower hypothesis, where the evolutionary maintenance of costly polyandry cannot be considered in isolation from the very strong selection that acts against virgin death [5, 9, 10].

We first discuss the seemingly contradictory results of low additive genetic (co)variance observed in the quantitative genetic experiment and the (correlated) response to artificial selection. Firstly, lack of statistical support for additive genetic (co)variance does not mean that there is no genetic (co)variance, only that we were unable to detect it with the statistical power that we had. Indeed, very low additive genetic variance in latency to mate would align with strong selection depleting genetic variation in this trait, supporting the spectre of the Wallflower hypothesis. Secondly, for a trait under strong selection, we might expect genetic variation to be caused by rare alleles and/or alleles with nonadditive effects. Notably, in the quantitative genetic experiment, we detected substantial dam variance. Within our design, we could not partition the contribution of maternal effects from dominance (among others) effects, and so dominance genetic variation may have contributed to that variance component. Interestingly, the presence of dominance variance may have facilitated the artificial selection response if nonadditive genetic variance was converted to additive genetic variation, though the impact of any such conversion is unclear [37]. Additionally, if genetic variation was largely caused by rare alleles that increased mating latency, then we should see an asymmetrical response to selection (only observed in the direction of increasing mating latency) that accelerates after an initial lag [38]. Our observed selection response was not inconsistent with the latter part of this theory, however, in terms of symmetry of the response, it would be very challenging to shorten mating latency in this population. Thus, while we do not know the underlying genetic architecture of female latency to (re)mate and, therefore, cannot test these questions directly, we stress that our results are not contradictory, but instead align with the Wallflower hypothesis.

An analysis of divergent populations, particularly those differing in mate encounter rates, could provide an instructive test of the hypothesis that genetic variation in virgin mating speed is eroded by selection against virgin death. We might expect populations to be fixed for different collections of alleles that affect female mating and remating, thus even though V_*A*_ *within* those populations might be largely absent (such is the case for desiccation resistance in rainforest *Drosophila* [39]), there might well be among-population covariance in virgin mating and the probability of remating as a non-virgin. Evidence consistent with different patterns among *versus* within populations comes from *Callosobruchus chinensis* in which a comparison across 10 strains of the first mating and subsequent remating rate of females was indeed moderately strong and in the expected direction (*r*_S_ = 0.55, *N* = 10, *P* = 0.098 [40]). This contrasts with a single population of these beetles subjected to divergent selection on remating rate, where there was no evidence of a correlated response in virgin mating [41].

How do the low levels of additive genetic variation in virgin matings square with the fact that there is no shortage of studies revealing heritable variation in polyandry [18-25] and responses to selection on polyandry [41-44]? We focussed our selection experiment on trying to increase the mating latency of virgins, as the Wallflower hypothesis explicitly predicts reluctance to mate at this stage to have carry-over effects to *subsequent* matings [5, 9, 10]. The reverse, (selecting on remating probability and examining virgin mating rates) has generated evolutionary responses in remating rates, but failed to change the virgin mating rate in either *Callosobruchus chinensis* [41], *Drosophila melanogaster* [44] or *Nasonia vitripennis* [42]. The heritable variation in, and evolution of, non-virgin choice that appears to be present more broadly may be facilitated by the novel environment (as compared to the ‘virgin environment’) that are delivered through the ejaculate of the first male and are not present in virgins. It is possible therefore that alleles that alter mating preferences can readily evolve if they are expressed in response to the numerous environmental effects in the post-mating environment (e.g., male seminal fluid constituents). This could decouple virgin and non-virgin mating behaviour (‘plasticity’ in Kokko and Mappes [5] terminology) and allow divergent mating acceptance thresholds in virgins and non-virgins. Nevertheless, this kind of decoupling is unidirectional; any allelic variation that is not environment specific but is generic to virgin and non-virgin mating will be rapidly eroded because of the costs of prolonging virginity. If this is the case, the Wallflower hypothesis cannot be adequately tested by selection on non-virgin mating frequency and its correlated effects.

Dam variance explained a high proportion of the variance in the quantitative genetic experiment. As noted above, within our design, we could not separate dam variance from dominance or other common environmental effects (*e*.*g*., microclimatic variation). Maternal influences are well documented in a diversity of traits in seed beetles [45-49] and in the remating behaviour of the parasitoid wasp *Nasonia vitripennis* [22]. Dominance genetic variation for female remating behaviour has been documented in fruit flies and house flies [50-52] and *Callosobruchus chinensis* [40]. Female size might be a candidate trait contributing to this variance component, although size does not correlate with latency to mate in this population (phenotypic correlation r_1880_ = -0.02, P = 0.3 Wilson and Tomkins Unpublished). Decomposing the dam variance would be of value for understanding variation in female mating latency, but we could not do that here.

If, as our results suggest, the Wallflower effect increases female responsiveness and thereby erodes the potential for female choosiness in the first mating, this alone is important because it weakens pre-copulatory sexual selection, particularly where there is not strong last-male sperm precedence [5, 10]. This weaker selection on males may help to maintain genetic variation, and can undermine important co-evolutionary processes such as the linkage between traits and preferences [9]. We note that our measure of ‘choosiness’ is female latency to mate, which does not explicitly test female discriminatory prowess; rather, her responsiveness to mate [53]. In the quantitative genetic experiment, half of the virgin females mated within 26 seconds (immediately on contact with the male) and 82% mated within a minute. Such rapid mating is suggestive of the fast and indiscriminate mating that would be consistent with the premise of the Wallflower effect.

Importantly, we found that females selected to be slow to mate as virgins, were even slower to remate than the fast-mating selected lines, and, than expected from their virgin mating latency, as captured in the selection history by mating status interaction (Fig. 1C). This effect illustrates both the problem and a possible solution to the risk of virgin death. It illustrates the problem, because it shows that selection on virgin mating latency carries over onto subsequent mating decisions and therefore that the wallflower effect has currency; it illustrates a potential solution, because it reveals that selection for virgin mating latency, increased mating status dependent plasticity. Importantly, this is not the plasticity envisaged as the ability to mate indiscriminately and then ‘trade-up’ by restoring choice: we found no evidence for this in the fast selected lines (Fig. 1, B&C). Rather, this is evidence for an effect whereby if females express some level of virgin discrimination, that there can thereafter be a heightened non-virgin discrimination– potentially increasing female fitness due to this form of ‘trading-up’. Thus, on the assumption that responsiveness and discrimination are correlated, our results highlight that non-virgin discrimination is dependent on the presence of some level of virgin discrimination, and the cost of virgin death constrains the evolution of non-virgin discrimination and trading up, increasing rates of polyandry. These results illustrate how mate encounter rates and sperm precedence patterns are likely key to where the evolutionary balance lies between avoiding the catastrophic fitness cost of virgin death and expressing mate discrimination as either a virgin or a nonvirgin [5, 10].

Interestingly, we recently documented genetic covariance in the choosiness of virgin and non-virgin *Drosophila melanogaster* (Dugand *et al* unpublished). That finding suggests that virgins and non-virgins align in their perceptions of male quality and further cautions against thinking that plasticity allows females free reign to ‘trade up’ after an indiscriminate first mating. Rather, together with the data presented here, it fuels the notion that the avoidance of virgin death may impact choosiness in females throughout their lives, and that the coevolution of choosiness and male trait expression (and all that goes with it), might indeed be affected [9]. In other words, if selection causes virgin females to mate quickly, genetic correlations across virgin and non-virgin status will have critically important implications whether virgin females are discriminatory or not. A key nuance of the Wallflower hypothesis is the preference-dependence of virgin death, i.e. that some females might be so unresponsive to male courtship that they die without mating at all. Selection for virgin death would be hard to execute; nevertheless, we have shown that within lineages experiencing selection for being less responsive to male courtship, those individuals that are less responsive to male courtship are less likely to mate polyandrously. This means that a female’s responsiveness to male courtship as a virgin does indeed affect polyandry. By enriching genetic variance for unresponsiveness to male courtship, we revealed the latent driver of the Wallflower hypothesis.

## Conclusions

The evidence that we have gathered is consistent with the premise and predictions of the wallflower effect; first, that there is extremely strong selection to mate at least once, consistent with an erosion of additive genetic variation for mating behaviour, and second, that by selecting on virgin mating latency we have gathered evidence for alleles affecting the readiness of virgin females to mate, that have carry-over effects on the probability of them subsequently remating. Our conclusion is that the wallflower hypothesis is a plausible evolutionary model for the scarcity of monandry [5]. Furthermore, although this work is couched in terms of understanding costly polyandry, it serves to explain beneficial polyandry as well [5]. For example, if we imagine an evolutionary experiment where lineages of females were forced into initially costly polyandry - we would expect some to adapt to the polyandry, reducing its harm or benefiting from it [6, 54]. Hence if the Wallflower effect enforces polyandry, this alone can explain the origins of both beneficial and costly fitness consequences that arise [5, 54].

## Acknowledgements

We thank the members of the Centre for Evolutionary Biology for feedback. This Research was funded by an Australian Research Council (FT110100500, DP130100783 & DP210100868).

## Extended Data

**Extended Data Table 1.**
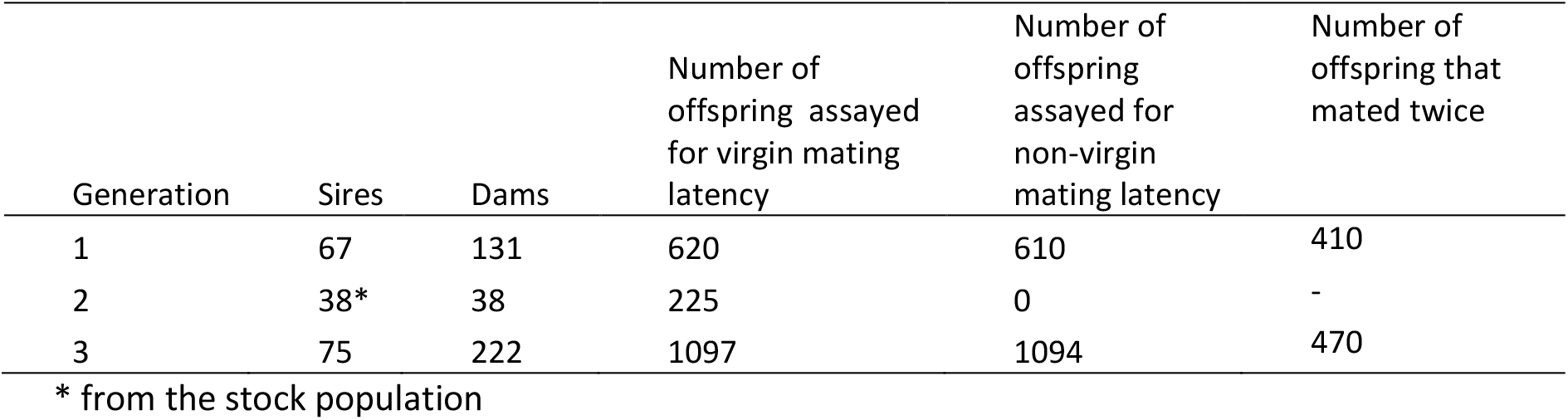
Pedigree structure and number of offspring assayed.

| Generation | Sires | Dams | Number of offspring assayed for virgin mating latency | Number of offspring assayed for non-virgin mating latency | Number of offspring that mated twice |
| --- | --- | --- | --- | --- | --- |
| 1 | 67 | 131 | 620 | 610 | 410 |
| 2 | 38* | 38 | 225 | 0 | - |
| 3 | 75 | 222 | 1097 | 1094 | 470 |

\* from the stock population

**Extended Data Table 2.**
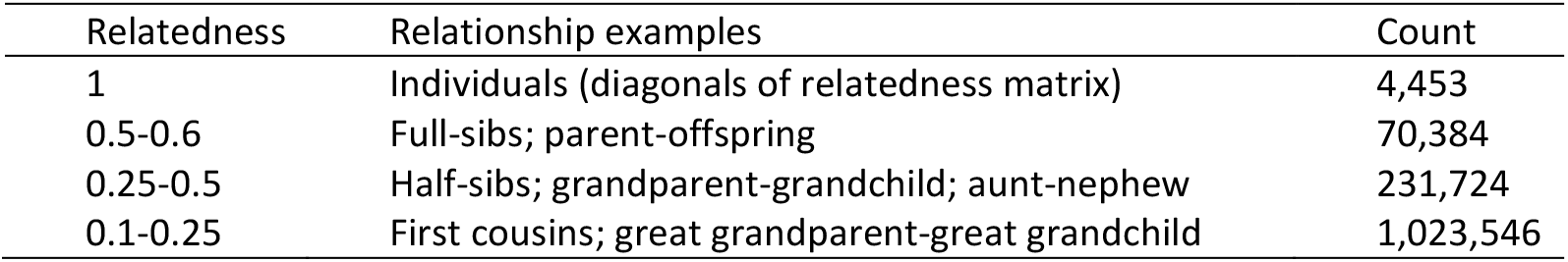
Summary of the relationships within the additive genetic relatedness matrix.

| Relatedness | Relationship examples | Count |
| --- | --- | --- |
| 1 | Individuals (diagonals of relatedness matrix) | 4,453 |
| 0.5-0.6 | Full-sibs; parent-offspring | 70,384 |
| 0.25-0.5 | Half-sibs; grandparent-grandchild; aunt-nephew | 231,724 |
| 0.1-0.25 | First cousins; great grandparent-great grandchild | 1,023,546 |

**Extended Data Table 3.** Estimates of the variances, the proportion of variance explained and the correlations for each of the group parameters from the quantitative genetic animal model of female latency to mate during the first and second mating. Estimates are the posterior means, with 95 % compatibility intervals shown in parentheses.

| Group parameter | Variance (first mating) | Proportion of total variance (first mating) | Variance (second mating) | Proportion of total variance (second mating) | Correlation |
| --- | --- | --- | --- | --- | --- |
| additive genetic | 0.033 [0.000, 0.130] | 0.208 [0.000, 0.706] | 0.066 [0.000, 0.336] | 0.052 [0.000, 0.255] | -0.041 [-0.601, 0.548] |
| dam | 0.037 [0.000, 0.103] | 0.250 [0.001, 0.710] | 0.128 [0.003, 0.300] | 0.107 [0.003, 0.280] | -0.233 [-0.712, 0.380] |
| date | 0.084 [0.009, 0.369] | 0.393 [0.077, 0.837] | 0.616 [0.112, 2.516] | 0.377 [0.128, 0.769] | NA |
| residual | 0.022 [0.000, 0.081] | 0.149 [0.000, 0.594] | 0.553 [0.338, 0.757] | 0.464 [0.150, 0.742] | -0.093 [-0.597, 0.468] |

**Extended Data Table 4.** Estimates of the variances and the proportion of variance for each of the group parameters from the quantitative genetic animal model of female receptivity during the second mating. Estimates are the posterior means, with 95 % compatibility intervals shown in parentheses.

| Group parameter | Variance | Proportion of total variance |
| --- | --- | --- |
| additive genetic | 0.106 [0.000, 0.631] | 0.101 [0.000, 0.478] |
| dam | 0.332 [0.107, 0.591] | 0.415 [0.135, 0.695] |
| date | 0.421 [0.138, 1.085] | 0.484 [0.204, 0.799] |

**Extended Data Table 5.** Effects of generation, selection treatment, and their interaction on latency to mate of virgin females.

| Effect | $\chi^2$ | d.f. | *P* |
| --- | --- | --- | --- |
| Generation | 0.00 | 1 | 1.000 |
| Treatment | 1.26 | 1 | 0.262 |
| Generation-by-treatment | 3.25 | 1 | 0.071 |

## Extended Data Figures

**Extended Data Figure 1.**
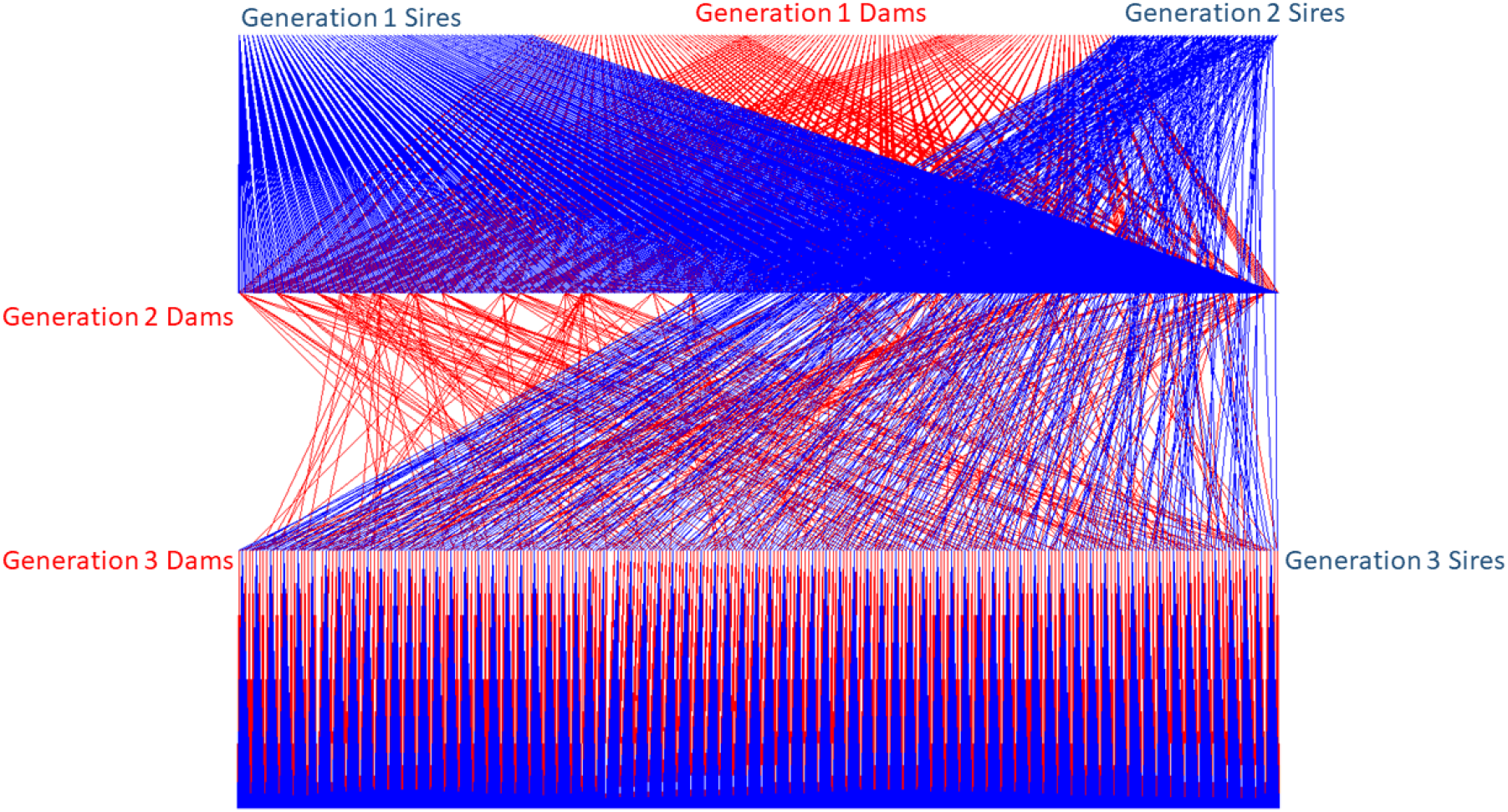
Halfsibling pedigree showing the introduction of stock population sires in generation 2 to pedigreed dams. Offspring from generations 1 and 3 were assayed for both first and second matings.

**Extended Data Figure 2.**
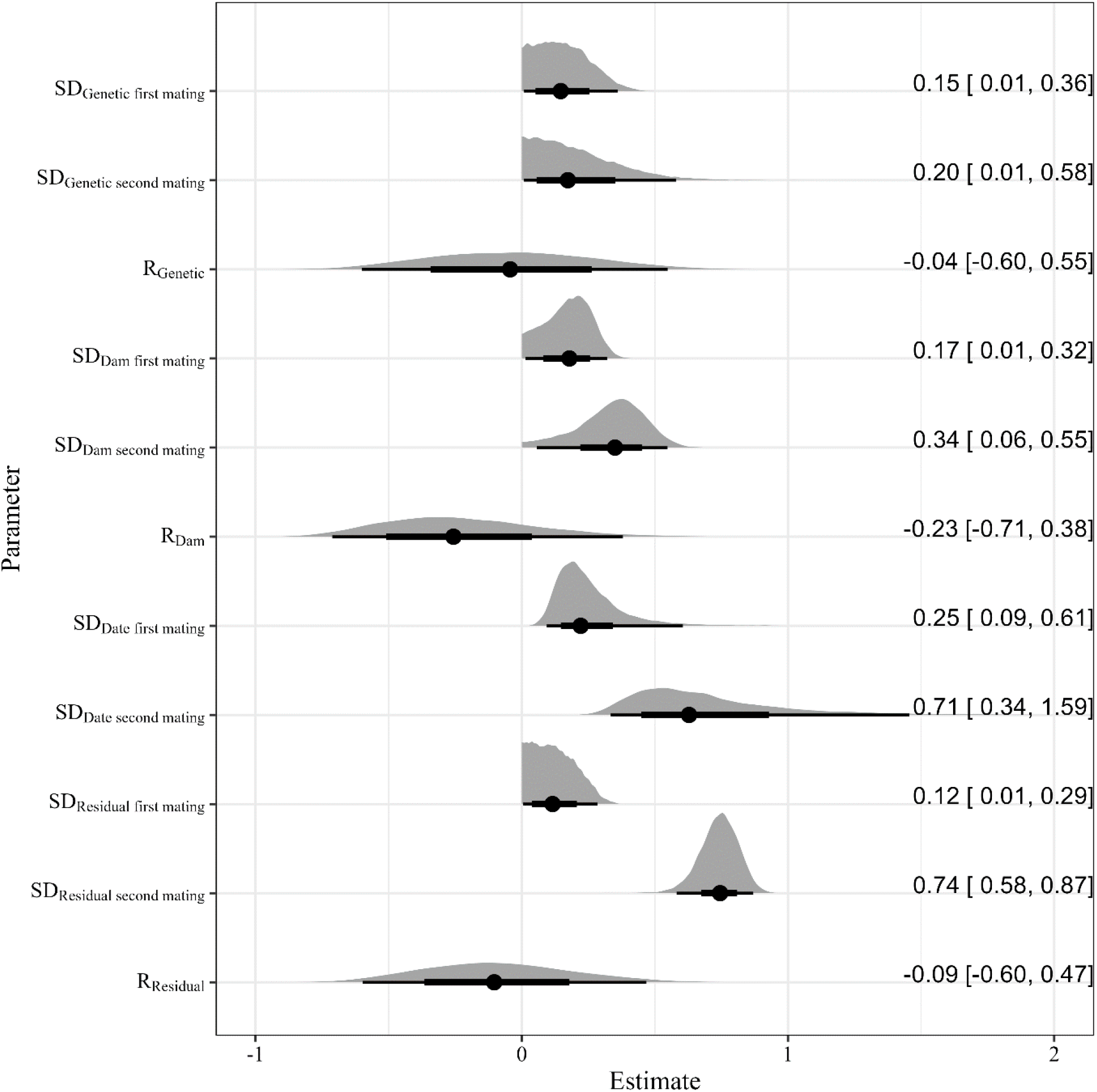
Posterior distributions for the group parameters estimated from the quantitative genetic animal model of female latency to mate during the first and second mating. SD = standard deviation, R = correlation. Density plots show the full posterior distribution, the points show the posterior mean, the thick whiskers the 50 % compatibility intervals, and the thin whiskers the 95 % compatibility intervals. The values for the posterior means are shown on the right of the figure, followed by the values for the 95% compatibility intervals in parentheses.

**Extended Data Figure 3.**
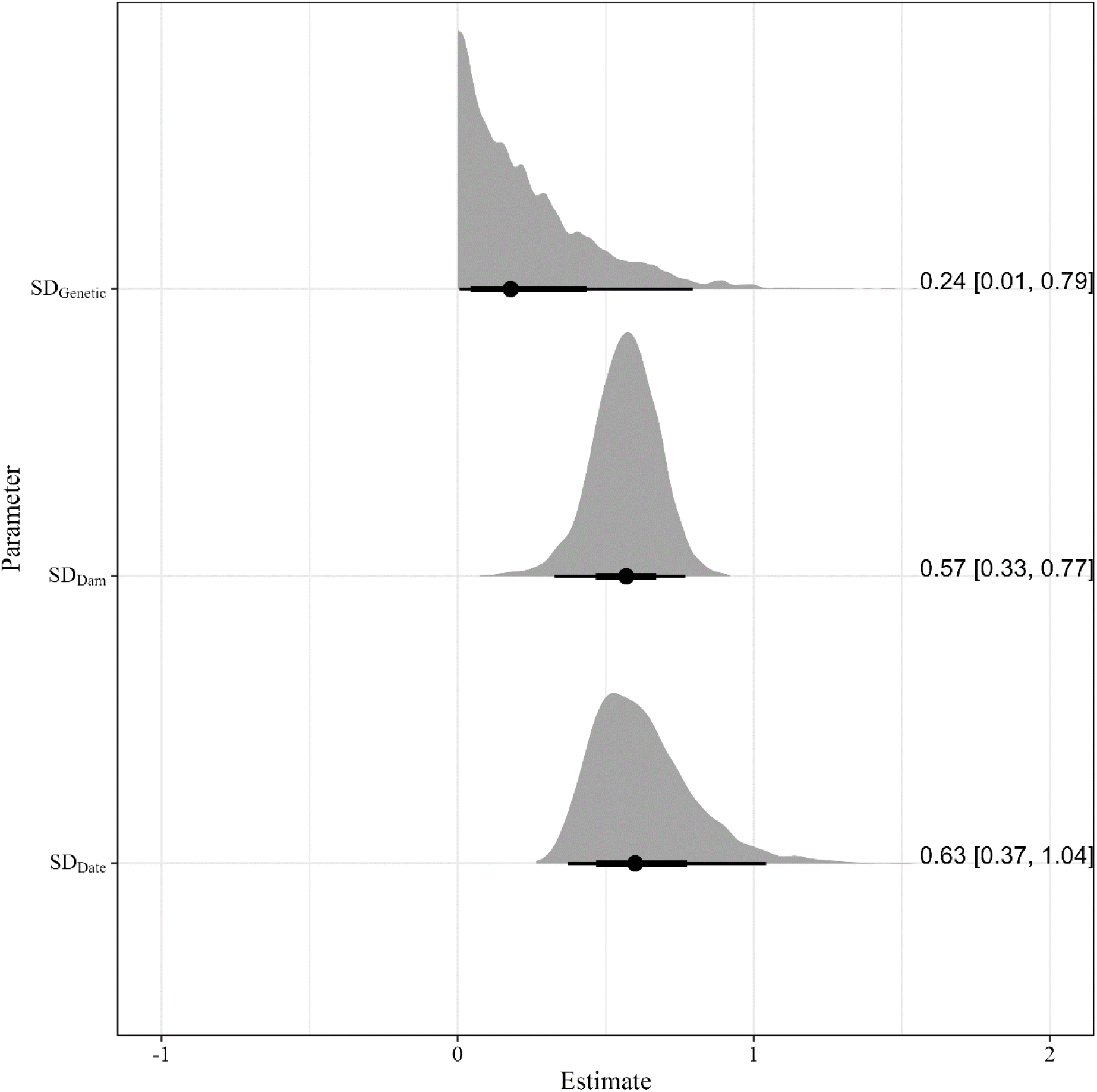
Posterior distributions for the group parameters estimated from the quantitative genetic animal model of female receptivity during the second mating. SD = standard deviation. Density plots show the full posterior distribution, the points show the posterior mean, the thick whiskers the 50 % compatibility intervals, and the thin whiskers the 95 % compatibility intervals. The values for the posterior means are shown on the right of the figure, followed by the values for the 95% compatibility intervals in parentheses.

## Notes

### Competing Interest Statement

The authors have declared no competing interest.

